# Agricultural intensification reduces microbial network complexity and the abundance of keystone taxa in roots

**DOI:** 10.1101/416271

**Authors:** Samiran Banerjee, Florian Walder, Lucie Büchi, Marcel Meyer, Alain Y. Held, Andreas Gattinger, Thomas Keller, Raphael Charles, Marcel G.A. van der Heijden

**Affiliations:** Agroscope, Department of Agroecology & Environment, Reckenholzstrasse 191, 8046 Zürich, Switzerland; Agroscope, Plant Production Systems, Route de Duillier 50, 1260 Nyon, Switzerland; Natural Resources Institute, University of Greenwich, United Kingdom; Research Institute of Organic Agriculture FiBL, 5070 Frick, Switzerland; Swedish University of Agricultural Sciences, Department of Soil & Environment, Box 7014, 75007 Uppsala, Sweden; Research Institute of Organic Agriculture FiBL, Jordils 3, 1001 Lausanne, Switzerland; Department of Plant and Microbial Biology, University of Zürich, Zürich 8057, Switzerland; Institute of Environmental Biology, Utrecht University, 3508 TC, Utrecht, The Netherlands

**Author notes:** Authors contributed equally to this work Correspondence.

## Abstract

Root-associated microbes play a key role in plant performance and productivity, making them important players in agroecosystems. So far, very few studies have assessed the impact of different farming systems on the root microbiota and it is still unclear whether agricultural intensification influences network complexity of microbial communities. We investigated the impact of conventional, no-till and organic farming on wheat root fungal communities using *PacBio SMRT sequencing* on samples collected from 60 farmlands in Switzerland. Organic farming harboured a much more complex fungal network than conventional and no-till farming systems. The abundance of keystone taxa was the highest under organic farming where agricultural intensification was the lowest. The occurrence of keystone taxa was best explained by soil phosphorus levels, bulk density, pH and mycorrhizal colonization. The majority of keystone taxa are known to form arbuscular mycorrhizal associations with plants and belong to the orders *Glomerales*, *Paraglomerales*, and *Diversisporales*. Supporting this, the abundance of mycorrhizal fungi in roots and soils was also significantly higher under organic farming. To our knowledge, this is the first study to report mycorrhizal keystone taxa for agroecosystems, and we demonstrate that agricultural intensification reduces network complexity and the abundance of keystone taxa in the root microbiota.

## Introduction

Agricultural intensification is one of the most pervasive problems of the 21st century [1]. To keep pace with the ever-increasing human population, the total area of cultivated land worldwide has increased over 500% in the last five decades [2] with a 700% increase in the fertilizer use and a several-fold increase in pesticide use [3, 4]. Agricultural intensification has raised a wide range of environmental concerns, including poor nutrient-use efficiency, enhanced greenhouse gas emissions, groundwater eutrophication, degradation of soil quality, and soil erosion [4, 5]. Alternate farming systems such as conservation agriculture (e.g., no-till) and organic farming have been widely adopted to reduce such adverse environmental effects [6, 7]. Organic farmlands represent 2.5% of the total arable lands in Europe, and over 3.5% in Switzerland [8], although they may reduce yield and yield stability [9]. The adoption of no-till globally has increased by approximately 233% in the last decade and it is over 3% of the total arable lands in Switzerland [10]. These farming systems are adopted to maintain environmental sustainability and ecosystems services, and at the heart of ecosystem services lies the contribution of microbial communities [11–13].

Microbial communities play an indispensable role in ecosystems and render a wide range of services [12, 14–16]. In agroecosystems, microbes modulate a number of processes, including nutrient cycling, organic matter decomposition, soil aggregate stabilization, symbiotic and pathogenic interactions with plants, and thereby play an essential role in the productivity and sustainability of agroecosystems [5, 12, 17]. The agricultural intensity with high resource-use and low crop diversity can affect soil-and plant-associated microbiota, with subsequent impact on ecosystem services [18, 19]. Increasing adoption of no-till and organic farming also warrants an investigation of their effects on microbial communities.

Previous studies comparing the effects of conventional, no-till and organic farming have mostly focused on the soil microbiome [20–22], and our understanding of the impact of these farming systems on root-associated microbiota is minimal.

Root-associated microbiota plays a key role in determining the above-ground productivity [23, 24]. No-till farming may affect root architecture and root distribution in soil, with a subsequent effect on microbial recruitment into the roots [25]. However, very few studies have assessed the effect of no-tillage on root microbial communities, and the ones that investigated root microbiota have only focused on root bacteria [26] or specific fungal groups, including arbuscular mycorrhizal fungi (AMF) using traditional techniques [27, 28]. Furthermore, the impact of agricultural intensification on the overall root fungal communities is still poorly understood [29, 30]. Plant root harbours a diverse assemblage of endophytic fungi that form symbiotic, parasitic or pathogenic associations, and through such associations, play a key role in plant diversity, community composition and performance [31]. The widespread symbiosis of AMF and the array of benefits rendered by these fungi are now well-established [32, 33]. Moreover, mycorrhiza like endophytes, *Piriformospra indica*, also promote plant growth, stress tolerance and induce local and systemic resistance to pathogens [34]. *Trichoderma* spp. have also been shown to enhance plant growth and systemic resistance to plant pathogens [35]. Thus, the structure and composition of root fungal communities play an important role in agroecosystems, and yet the effect of agricultural intensification on root fungal communities remains poorly understood.

The structure of a microbiome has substantial effects on its functioning [36]. However, studying the structure of a microbiome is not simple mainly due to complex interrelationships among the myriad of members. Microbial co-occurrence networks can unravel such relationships and offer insight into community structure [37, 38]. Network analysis has been found particularly useful in recent years to understand how microbe-microbe associations change in response to environmental parameters [39–42]. Network scores can also be used to statistically identify the keystone taxa, i.e. taxa that have a large influence in the community [43, 44]. Recent studies have demonstrated that such highly connected taxa can explain microbiome compositional turnover better than all taxa combined [45]. It has also been observed that the impact of abiotic factors and host genotypes on the plant microbiome is facilitated via keystone taxa [46], and the root microbial network complexity is linked to plant survival [47]. Agricultural intensification may alter the structure of root microbial network and the abundance of keystone taxa, which in turn may have implications for crop performance [48]. However, so far, it has not been investigated whether root microbial networks differ between organic, conservation and conventional agriculture. A pertinent question is whether mycorrhizal fungi that are widely regarded for their role in plant productivity can also act as keystone taxa in the microbial community.

Here we explored the impact of farming systems on the fungal community structure using the latest *PacBio SMRT sequencing* and network analysis of wheat root samples collected from 60 farmlands in Switzerland. We aimed to address the following questions: a) does agricultural intensity affect the structure and composition of wheat-root fungal communities? b) do network complexity and the abundance of keystone taxa vary between conventional, no-till and organic farming? c) which taxa act as keystone and what are the drivers of such taxa in the root microbiota?

## Material and methods

### Site selection and sampling

Soil samples were collected in early May 2016 from wheat fields in 60 agricultural farmlands in the northeast and southwest regions of Switzerland (Figure S1). Wheat fields were either managed conventionally with tillage, conventionally under no-tillage, or organically under a mouldboard plough tillage. Farming systems were distributed equally in both regions, and each system was represented by 20 farmlands. At each farmland, 18 soil cores (4 cm diameter) were collected at 0-20 cm depth with a hand auger (Figure S2). These 18 samples were mixed and pooled to obtain a representative sample for a farm. The auger was cleaned between sites. Five undisturbed cylindrical soil cores of 100 ml volume and 5.1 cm diameter were collected for bulk density measurement and the median of the five measures was considered as the estimate of bulk density for each field. Root samples were collected in June 2016. At each site, ten wheat plants, five per transect, were excavated using a fork spade. Shoots were cut off at the height of approx. 5 cm and all roots of a specific site were pooled in a plastic bag for subsequent processing. Samples were placed on ice in a cooler box for transfer to the laboratory. Soil samples were processed on the same day as the collection by removing plant materials, homogenizing and passing through a 2 mm sieve. Sub-samples were taken for various soil physicochemical and biological analyses and stored at appropriate temperature as required.

### Plant and soil analyses

In the lab, roots were thoroughly cleaned under cold tap water. Subsequently, fine roots (< 1 mm) were cut into small pieces of about 1 cm length and thoroughly mixed. A subsample of 2 g of fine roots was stored in 1.5 Eppendorf tubes, lyophilized and stored at -20°C for DNA extraction. The rest of the samples was used to determine AMF colonization by estimating the abundance of arbuscules, hyphae or vesicles according to a modified line intersection method [49] using a minimum 100 intersections per slide of two technical replicates and applying a blind procedure throughout the quantification process to avoid subjectivity related to the origin of the sample. Total phosphorus (P), plant available P, pH, and bulk density were measured using the Swiss standard protocols (FAL, 1996). Plant available P was measured according to Olsen *et al.* (1954). The abundance of AMF in soil was assessed by the phospholipid fatty acid (PLFA) analysis [52]. We quantified the abundance of AMF in soil by using the PLFA 16:1ω5, which is well-regarded as a biomarker for AMF because it constitutes a large proportion of total PLFAs in AMF, and strong correlations between AMF abundance in the soil and concentrations of the PLFA 16:1ω5 have been observed previously [53].

### DNA extraction and SMRT sequencing

For each sample, 200 mg of roots (dry weight) was used for DNA extraction using 600 mL of Nucleo spin lysis buffer PL1 for 15 min at 65 °C followed by the NucleoSpin Plant II kit (Macherey & Nagel, Düren, Germany). The DNA samples were amplified with the primer pair *ITS1F-ITS4* [54, 55] targeting the entire ITS region (approx. 630 bp) [56]. The forward and reverse primers were synthesized with a 5-nucleotide-long padding sequence followed by barcode tags at the 5’ end to allow multiplexing of samples within a single sequencing run [57]. Library preparation and SMRT sequencing were conducted at the Functional Genomics Centre Zurich (http://www.fgcz.ch) on the PacBio® RS II Instrument (PacBio, San Diego, CA, USA). Details of PCR conditions and sequence data processing are described in the *Supplementary Information*. In brief, the SMRT Portal was used to extract the circular consensus sequences (CCS) from the raw data (available from the European Nucleotide Archive, study accession number: PRJEB27781). The CCS of at least five passes yield similar error rates as 454 or MiSeq sequencing platforms [56, 57]. The CCS reads were quality filtered in Mothur (v.1.35.0) [58]. Quality reads were demultiplexed based on the barcode-primer sequences using *flexbar* [59]. *De novo* chimera detection was performed on quality reads using UCHIME [60]. To avoid unwanted multi-primer artefacts, we deleted reads where full-length sequencing primer was detected within the read [61]. We clustered the quality sequences into operational taxonomic units (OTUs) at ≥ 98% sequence similarity with the UPARSE series of scripts [62]. Reads were de-replicated, and single-count and chimeric sequences were excluded for OTU delineation. The OTUs of low abundance (less than 0.1% global abundance and less than 0.5% abundance within a specific sample) were removed from the dataset (Figure S3). On average 357 OTUs were found per site and a total of 823 OTUs for all 60 sites. The OTUs were classified taxonomically against the UNITE database [63]. The OTU and taxonomy tables were filtered to exclude OTUs classified as non-fungal.

### Statistical analyses

Alpha diversity indices such as OTU richness, Sheldon evenness and Shannon-Weaver index were calculated from the rarefied fungal OTU table using the *phyloseq* package [64] in R v3.4 [65]. The effect of farming systems and wheat varieties on fungal community structure was assessed by performing PERMANOVA and canonical analysis of principal coordinates (CAP) with 999 permutations in PRIMER-E (PRIMER-E, Plymouth, UK). Fungal beta diversity patterns were only assessed on OTUs that were present in at least two samples. Homogeneity of multivariate dispersions was checked with the PERMDISP test using the Bray-Curtis similarity matrix in PRIMER. We also identified the indicator taxa for each farming system using the *indicspecies* package in R [66]. Co-occurrence patterns in fungal communities were assessed by performing network analysis using the maximal information coefficient (MIC) scores in MINE statistics [67]. MIC is an insightful score that reveals positive, negative and non-linear associations among OTUs. To minimize pairwise comparisons, network analysis was performed on OTUs that were present in at least two samples, resulting in 826 fungal OTUs. The overall meta-network was constructed with 60 samples whereas the three farming specific networks were constructed with 20 samples each. The MIC associations were corrected for false discovery rate (FDR) [68] and the final networks were constructed with relationships that were statistically significant (P<0.05) after FDR correction. The networks were then visualized in Cytoscape version 3.4.0 [69]. The *NetworkAnalyzer* tool was used to calculate network topology parameters. We also evaluated networks against their randomized versions using the Barabasi-Albert model available in *Randomnetworks* plugin in Cytoscape v2.6.1. The structural attributes of fungal networks such as degree distribution, mean shortest path, clustering coefficient were different from random networks with an equal number of nodes and edges. The OTUs with the highest degree and highest closeness centrality, and the lowest betweenness centrality scores were considered as the keystone taxa [43]. We calculated the influence of various taxa in network stability by dividing the number of nodes belonging to a particular taxon by the number of connections (edges) it shared. Finally, we performed Random Forest Analysis [70] to explore the determinants of the identified keystone taxa with 999 permutations using the *randomforest* and *rfPermute* packages in R [71]. The best predictors were identified based on their importance using the *importance* and *varImpPlot* functions. Increase in node purity and mean squared error values were used to calculate the significance of the predictors using the *randomForestExplainer* package [72]. The factors significant at P<0.01 were selected as the predictors of keystone taxa. Agricultural intensity index was calculated according to a previous study [73] based on the information collected from farmers (Büchi et al., submitted). Agricultural intensity index was estimated using the information on three anthropogenic input factors: fertilizer use, pesticide use and the consumption of fuel for agricultural machineries. These factors were also included for assessing agricultural intensity in a previous study [74].

## Results

### Overall structure and co-occurrence

Farming systems significantly influenced the root fungal community structure with three distinct clusters for organic, conventional and no-tillage fields (Figure 1A). A PERMANOVA test confirmed the significant effect of farming systems (pseudo F = 1.42; P<0.05). However, alpha diversity indices and the overall taxonomic composition did not vary between the conventional, no-till and organic systems (Figure S4, S5). A non-significant PERMDISP test (F = 2.072; P = 0.202) indicated homogenous dispersions of samples across systems. Further, a pairwise comparison in PERMDISP revealed that there was no significant difference in dispersions between organic and conventional (F = 1.068; P = 0.372), and organic and no-till (F = 0.870; P = 0.435). We found no impact of wheat varieties on community structure and this was reinforced by a non-significant PERMANOVA test (Pseudo F = 0.972; P = 0.595) (Figure S6). However, geographical locations i.e., northeast and southwest regions had an impact on root fungal community structure (Figure S7). Indicator species analysis was performed to test which taxa are characteristic for each of the three farming systems. Root inhabiting *Trichoderma*, a member of *Hypocreales*, was the only indicator taxon for conventional farming system whereas seven fungal taxa (e.g., *Cyphellophora, Myrmecridium, Phaeosphaeria, Cadophora, Pyrenochaeta, Solicoccozyma*, and *Conocybe*) were the indicator taxa for no-till farming (Table S1). Six taxa of *Sordariales, Cantharellales*, and *Agaricales* were indicator taxa for organic farming with *Chaetomium* and *Psathyrella* as the only known genera.

**Figure 1.**
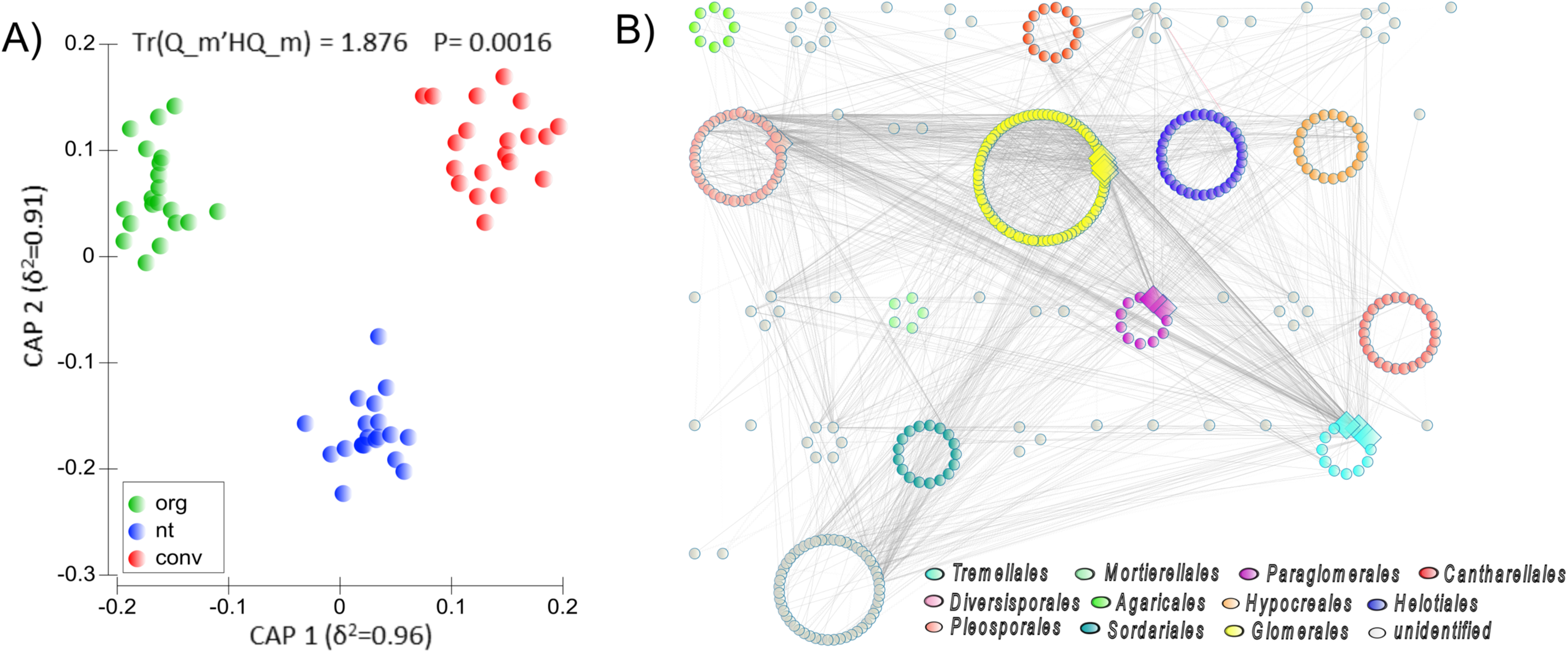
**A)** Canonical analysis of principal coordinates (CAP) revealing a significant impact of farming systems on fungal community structure. **B)** The overall network of root fungal communities across three farming systems. The overall network is arranged according to orders. White, red and wavy lines represent positive, negative and non-linear relationships, respectively. Large diamond nodes indicate the keystone taxa in the network. Top ten nodes with the highest degree, highest closeness centrality, and lowest betweenness centrality were selected as the keystone taxa [43]. Out of the ten keystone taxa in the overall network, seven belonged to mycorrhizal orders, *Glomerales, Paraglomerales*, and *Diversisporales*.

The overall network of root fungal communities in 60 samples revealed distinct co-occurrence patterns (Figure 1B). The meta-network consisted of 378 nodes and 1602 significant (P<0.05) edges. This network with strong power-law distribution of degrees had a diameter of 8, average number of neighbours of 8.476, and a clustering coefficient of 0.258. For the overall network, the top ten taxa with the highest degree, highest closeness centrality, and lowest betweenness centrality were selected as the keystone taxa (Table S2). Seven of these taxa belonged to arbuscular mycorrhizal orders *Glomerales, Paraglomerales*, and *Diversisporales* and the remaining three belonged to *Tremellales*. Indeed, the majority of the associations were from these four orders with *Glomerales* forming the largest guild with the maximum number of nodes and associations in the network. Overall, farming systems significantly affected fungal community structure with mycorrhizal orders playing a major role in the network complexity.

### Farming specific co-occurrence networks

Owing to the significant difference in fungal community structure across three farming systems, we further evaluated root fungal networks for each farming system separately. The networks displayed remarkable differences in their structure and topology (Figure 2). The network of conventional farming consisted of 261 nodes (e.g., taxa) and 315 edges (associations between taxa) while the no-till network consisted of 267 nodes and 341 edges. In stark contrast, the organic farming network consisted of 301 nodes and 643 edges. The average number of neighbours and the clustering coefficient of the organic farming network were also considerably higher than for the other two networks (Figure 2). The higher complexity and connectivity in the organic farming network were supported by the abundance of keystone taxa, taxa that are central to community structure. The organic farming network harboured 27 of such keystone taxa compared to two in the no-till network and none in the conventional one (Figure 2; Table S3). The majority of these keystone taxa belonged to the orders *Glomerales*, *Tremellales* and *Diversisporales* with a noticeable presence of taxa from the orders *Paraglomerales*, *Sebacinales* and *Hypocreales*.

**Figure 2.**
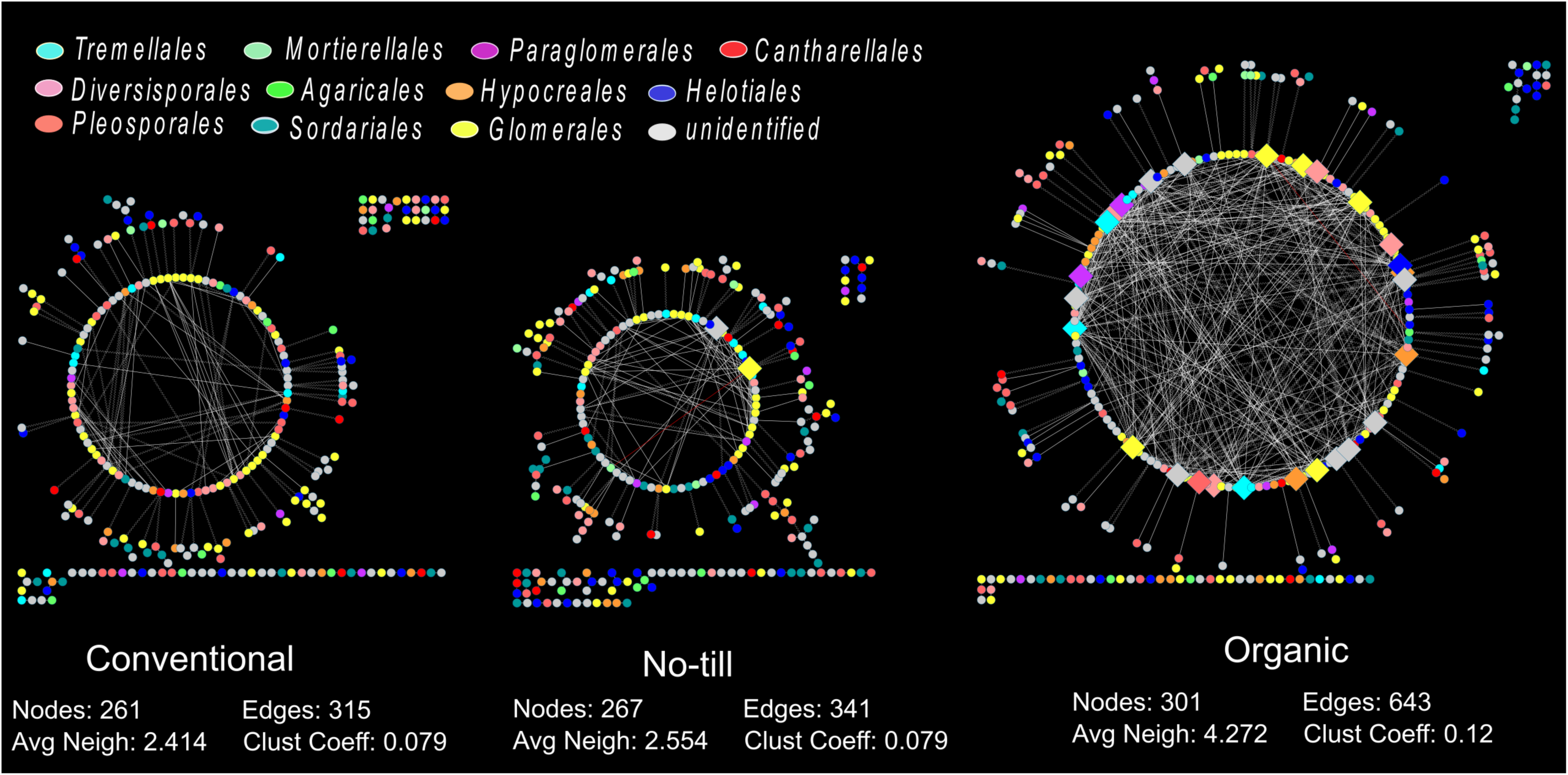
Farming system specific root fungal networks. Each network was generated with root samples from 20 farmlands belonging to that farming systems. Large diamond nodes indicate the keystone taxa whereas circular nodes indicate other taxa in the network. White, red and wavy lines represent positive, negative and non-linear relationships, respectively. Despite having similar number of nodes, the organic network displayed twice more edges and many highly connected nodes than no-till and conventional networks that were dominated by less connected peripheral nodes.

Higher connectivity in the organic farming network was visible in the distribution of degrees, which indicates the number of associations shared by each node in a network (Figure 3). The organic farming network had a much stronger power-law distribution than the conventional and no-till ones, despite the similar node distribution across root fungal orders (Figure S8). We calculated the proportional influence of various orders in the microbiota by dividing the number of nodes belonging to a particular order by the number of connections (edges) it shared. It revealed the orders that exhibited maximum connections across three farming systems and thereby influence the network structure. Various orders exhibited considerable differences in their proportional influence in the stability of root microbiota. Orders such as *Sordariales* and *Agaricales* showed a major influence in the conventional network structure, and *Sordariales, Cantharellales* and *Mortierellales* in the no-till network. In addition to *Tremellales and Hypocreales*, three mycorrhizal orders *Glomerales*, *Paraglomerales* and *Diversisporales* showed a major influence on network stability under organic farming. Overall, the organic farming network formed a much more complex network and harboured more keystone taxa than the other two farming networks.

**Figure 3.**
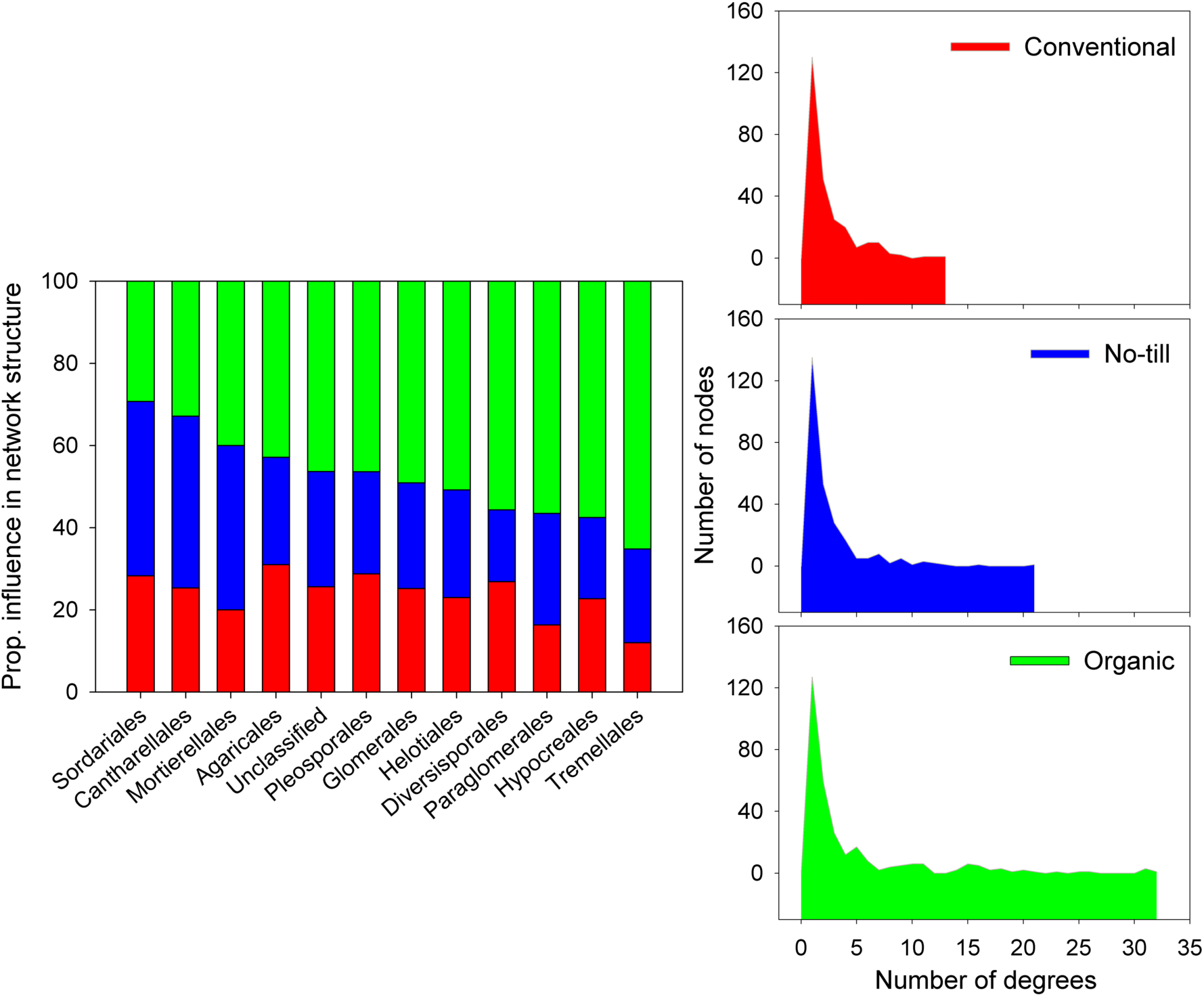
Proportional influence of various fungal orders in affecting the stability of root microbiota (left panel). The influence was calculated by diving the number of nodes belonging to a particular fungal order by the number of connections (edges) it shared. It illustrates the orders that exhibit maximum connections across farming systems and thus influences network structure most. Distribution of degrees in three farming systems (right panel with three plots). Degree indicates the number of associations shared by each node in a network. In conventional, farming, the number of degrees was limited to a maximum of 12 compared to the no-till network that had a maximum of 22 degrees. On the other hand, organic farming had many nodes with over 20 degrees.

### Drivers of keystone taxa

Random forest analysis revealed that soil phosphorus content, bulk density, pH, and mycorrhizal colonization best explained (P<0.01) the occurrence of keystone taxa (Figure 4). Most of these parameters were also significantly (P<0.05) correlated with the alpha-diversity indices, indicating their importance for the overall root fungal communities (Table S4). The majority of keystone taxa belonged to mycorrhizal orders, and mycorrhizal colonization of wheat roots was significantly (P<0.01) higher in the organic fields than in the conventional and no-till fields (Figure S9). Consistent with this, the abundance of mycorrhizal PLFA in soil were also significantly (P<0.01) higher in the organic fields. Agricultural intensity had a significantly negative impact on mycorrhizal colonization in roots and the abundance in soils (Figure 4). Agricultural intensity was significantly (P<0.05) different across three farming systems with conventional as the most intensive and organic as the least intensive system (Figure S9). This trend was opposite for the abundance of keystone taxa i.e., the number of keystone taxa was 27 in the organic farming network, 2 in the no-till, and 0 in the conventional network. Taken together, the root fungal network complexity, abundance of keystone taxa and mycorrhizal abundance showed an opposite trend to that of agricultural intensification across farming systems.

**Figure 4.**
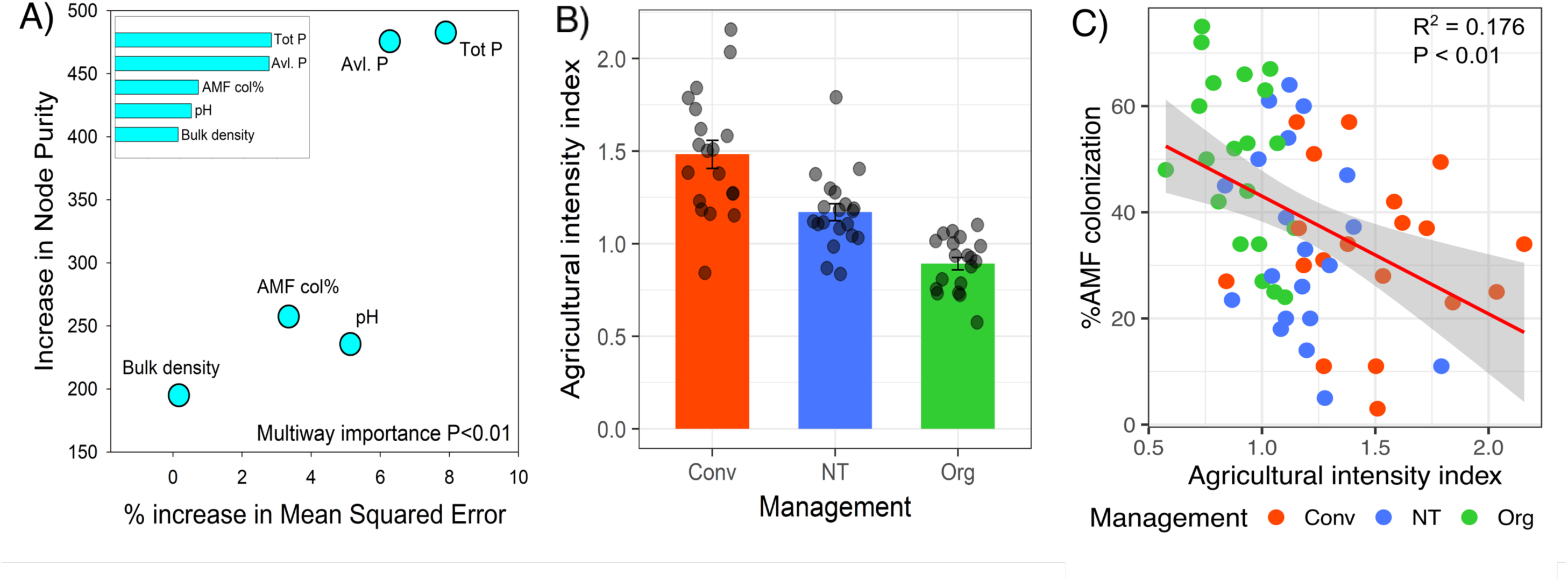
A) Results of Random Forest Analysis showing the relative contribution of various factors in determining the abundance of keystone taxa. The mean squared error (MSE) indicates the prediction accuracy of each factor. The top (P<0.01) five drivers were total phosphorus, plant available phosphorus (Olsen P), AMF root colonization, pH and bulk density. B) Agricultural intensity index across three farming systems. Agricultural intensity index was estimated using the information on three anthropogenic input factors: fertilizer use, pesticide use and the consumption of fuel for agricultural machineries. C) Relationship between agricultural intensity and mycorrhizal root colonization. Agricultural intensification had a significantly (P<0.01) negative impact on the root colonization of AMF. Agricultural intensity was the highest under the conventional farming and the lowest under the organic farming, which was opposite for the AMF colonization.

## Discussion

It is now well established that root-associated microbiota plays an important role in plant diversity, community composition and performance [24, 32, 75]. Consequently, it is important to understand how microbial communities harboured in crop roots are affected by farming systems and how key microbial players can be harnessed for ecological intensification of agroecosystems. However, with much of the previous work only focussing on the soil microbiota, our understanding of the effects of farming systems on root-associated microbiota is still rudimentary. Moreover, previous studies mostly focused on microbial alpha-and beta diversity patterns and the impact of different farming systems on microbial network structure is poorly understood.

Our results showed that agricultural intensity was the highest under conventional farming and the lowest under organic farming. The overall structure of root microbiota was significantly influenced by farming systems. This is also consistent with studies on soil microbiome where a large number of reports showed a significant impact of farming systems [20–22, 76, 77]. For the root microbiota, Hartman *et al.* (2018) observed in a farming experiment that root fungal communities were only affected by tillage intensity and not by conventional or organic farming. It should be noted that most of these studies investigated microbial communities in field-trials [20–22, 30, 77]. While a major strength of field-trials is that farming treatments are imposed under a homogenous management and at one location with a specific soil type, management effects on microbial patterns may be different in actual farmlands and thus, the results obtained at one location cannot be generalized. It is necessary to investigate whether microbial community characteristics observed in field-trials can be generalized in on-farm research and across many fields. To our knowledge, this is the first on-farm study on root fungal communities under different farming systems focussing on conventional, organic and conservation agriculture. Our results show that wheat roots under different farming systems harbour distinct fungal communities and with varying network complexity.

Microorganisms do not thrive in isolation and rather form complex association networks. Such networks hold special importance for gaining insight into microbiome structure and its response to environmental factors [37, 38, 46]. Our study highlights how farming systems impact the network structure of root microbiota, and uncovers, for the first time, that organic farming harbours a significantly more complex network than the conventional and no-till farming. The organic network exhibited a much stronger power-law distribution of degrees with many highly connected nodes whereas no-till and conventional networks were dominated by less connected peripheral nodes. It has been shown that complex networks with greater connectivity are more robust to environmental perturbations than simple networks with lower connectivity [78]. In this sense, the higher complexity of organic networks may indicate that the root microbiota under organic management is more resilient to environmental stresses as different taxa can complement each other and network complexity may provide an insurance for network stability if specific taxa go extinct. However, further studies are necessary to corroborate this observation.

Keystone taxa are the highly connected taxa that play important roles in the microbiome and their removal can cause significant changes in microbiome composition and functioning [43, 44]. Although previous studies have reported keystone taxa in various environments [40, 79, 80], reports on keystone taxa in the root microbiome are very limited. The organic farming network exhibited by far the highest connectivity and comprised most of the keystone taxa. It should be noted that fungal richness did not vary significantly between the farming systems nor did the node distribution, and yet we observed a clear difference in the network structure. Nodes in the organic network shared more associations and there were also many highly associated keystone taxa. The abundance of these keystone taxa did not vary between the three farming systems but these taxa shared considerably more associations in the organic farming (Figure S10). A previous study reported a gradient of root fungal assemblages from conventional to organic to natural grasslands, i.e., root microbiota in the organic farming was more developed than the conventional farming [81]. These observations indicate a possibility that microbiome complexity is not necessarily determined by the number of taxa in the community, but rather the number of associations that those taxa share amongst them.

Remarkably, the majority of these keystone taxa were arbuscular mycorrhizal fungi (AMF) belonging to the orders *Diversisporales, Glomerales*, and *Paraglomerales*. The symbiotic association of AMF that started more than 400 million years ago is formed by ∼80% of terrestrial plants [33, 82]. The observation that AMF can enhance plant productivity [83] make them a crucial player in agroecosystems. The importance of AMF for the root-associated microbiota, particularly under organic farming, is congruent with the higher abundance of AMF in roots and soils observed in the organic farmlands in this study (Figure S9). Mycorrhizal root colonization was significantly higher under organic farming than non-till and conventional farming systems. This pattern was also evident for mycorrhizal abundance in the soil as measured by AMF-specific PLFA concentration. While previous studies also found significantly higher AMF abundance and diversity in organic farmlands than in the conventional ones [81, 84], the pivotal role of AMF for the entire root-associated microbiota in agroecosystems is reported here for the first time.

One of the non-mycorrhizal keystone taxa in the organic farming belonged to the order *Sebacinales*. Members of this order are highly diverse root endophytes and are thought to form neutral and beneficial interactions with plants [85]. Our observation of *Sebacinales* as keystone taxa is consistent with a previous report that found a consistently higher abundance of *Sebacinales* in organic farmlands [29]. Since keystone taxa are linked to network complexity, beneficial endophytic keystone taxa such as AMF and *Sebacinales* may enhance the network complexity and thereby the stability of the root microbiome. Several other keystone taxa in the overall and organic networks belonged to the order *Tremellales*. This widespread group of Basidiomycetes contains many yeast species and have been reported in plant roots in temperate regions [86]. Members of this fungal order were also recently found as keystone taxa in the root microbiome across eight forest ecosystems in Japanese Archipelago [48]. Interestingly, we found that two of the keystone taxa (OTU_10, OTU_11) were members of the *Dioszegia* genus that was also found as keystone by Agler *et al.* (2016). It was shown that the effect of abiotic factors on microbiome was mediated via *Dioszegia* in *Arabidopsis thaliana*. The consistent identification of *Dioszegia* as a keystone taxon across studies suggests its importance and highlights a potential that it can be harnessed for manipulation of the plant microbiome. Future studies are now needed to specifically manipulate this taxon to test how it influences microbiome composition and functioning. There were no common fungal groups between indicator taxa and keystone taxa. It should be noted that indicator taxa are identified based on their higher/lower abundance under a particular farming system whereas keystone taxa are identified using a comprehensive algorithm that focuses on the number of associations a taxon shares and its position in the microbiome. Thus, indicator taxa and keystone taxa highlight different aspects of microbial communities.

Beyond statistical identification of keystone taxa, this study also revealed the factors driving their abundance in the wheat root microbiota. Bulk density plays a pivotal role in root elongation rates, rooting density and root architecture [87], which have significant implications for microbial recruitment into the roots. We found soil phosphorus levels and plant available phosphorus were the strongest determinants of keystone taxa. The majority of keystone taxa were mycorrhizal in nature, and phosphorus is well-acknowledged for its importance for mycorrhizal associations [88]. Similarly, soil pH is a known driver of fungal communities, especially, mycorrhizal fungi [89, 90]. Thus, the identification of soil phosphorus levels, pH and bulk density as the determinants of keystone taxa in root microbiota is plausible. The number of keystone taxa was the highest under organic farming where agricultural intensity was the lowest and conversely, intensification was the highest under conventional farming where root microbiota did not have any keystone taxa. Thus, agricultural intensification might have negatively affected the network complexity and keystone abundance in the root microbiota. It is important to mention that identification of keystone taxa are based on the analysis of correlations (associations) among taxa, and further research is necessary to show causality, in terms of the impact of keystone taxa on microbiome structure and functioning.

## Conclusions

The structure and composition of root microbiota play an important role in agroecosystems and yet there is a significant dearth of knowledge about the effect of agricultural intensification on the root microbiota. Our study shows that the microbiome complexity and the abundance of keystone taxa were the highest under organic farming where agricultural intensity was the lowest. The higher co-occurrence of members of microbial communities under the organic farming can be indicative of greater ecological balance and stability of the microbiome. A key strength of this study is that the samples were collected from 60 fields and the reported effects can be generalized because samples were taken from an extensive range of fields at different locations with different management regimes. The recent concept of smart farming (*sensu* Wolfert *et al.*, 2017) emphasizes *thinking outside the box*. The potential for harnessing plant microbiome for sustainable agriculture was also highlighted recently [92]. Mycorrhizal fungi are well-regarded for their effects on plant productivity and thus, mycorrhizal keystone taxa may be targeted as a tool for smart farming.

## Acknowledgement

We especially thank the farmers for allowing access to their farmlands and submitting survey reports. We also thank Julia Hess, Kexing Liu, Cindy Bally, Florent Georges, Thibault Granger, Marlies Sommer, Andreas Fliessbach and Raphael Wittwer for assistance with field-and lab work. This study was funded by the Swiss National Science Foundation (grant 31003A-166079) and National Research Program ‘Sustainable Use of Soil as a Resource’ (NRP 68) [grant 406840-161902].

## Author contributions

MvdH, RC and TK conceived and designed the study; FW, LC and MM conducted the sampling; AYH and MM performed molecular and mycorrhizal analyses; SB performed statistical analyses and wrote the manuscript; FW also contributed to manuscript preparation; all authors edited the manuscript.

## Sequence availability

All sequences generated in this study are available in the European Nucleotide Archive under the study accession number: PRJEB27781.

